# Ambiguous Body Ownership Experience Caused by Feeling–judgement Conflict: Evidence from Subjective Measurement in Rubber Hand Illusion using Integrated Information Theory

**DOI:** 10.1101/2021.11.29.470324

**Authors:** Takayuki Niizato, Yuta Nishiyama, Kotaro Sakamoto, Takumi Kazama, Tatsuya Okabayashi, Taiki Yamaguchi

## Abstract

Human body awareness is malleable and adaptive to changing contexts. The illusory sense of body-ownership has been studied since the publication of the rubber hand illusion, where ambiguous body ownership feeling, expressed as “the dummy hand is my hand even though that is not true”, was first defined. Phenomenologically, the ambiguous body ownership is attributed to a conflict between feeling and judgement; in other words, it characterises a discrepancy between first-person (i.e. bottom-up) and third-person (i.e. top-down) processes. Although Bayesian inference can explain this malleability of body image sufficiently, the theory does not provide a good illustration of why we have different experiences to the same stimuli – the difficulty lies in the uncertainty regarding the concept of judgement in their theory.

This study attempts to explain subjective experience during rubber hand illusions using integrated information theory (IIT). The integrated information Φ in IIT measures the difference between the entire system and its subsystems. This concept agrees with the phenomenological interpretation – that is, there is conflict between judgement and feeling. By analysing the seven nodes of a small body– brain system, we demonstrate that the integrity of the entire system during the illusion decreases with increasing integrity of its subsystems. These general tendencies agree well with many brain-image analyses and subjective reports; furthermore, we found that subjective ratings were associated with the Φs. Our result suggests that IIT can explain the general tendency of the sense of ownership illusions and individual differences in subjective experience during the illusions.

## 1 Introduction

The malleability of human body image shows that the perceptual system is not fixed but flexible. This malleability has been demonstrated in several experiments where the body awareness is altered depending on multimodal stimuli. One famous example is the rubber hand illusion (RHI) introduced in Botvinick and Cohen [1998], where subjects observe a brush stroking on a dummy hand while the subjects’ real hand, which is hidden behind a partition, is also stroked. Synchronized visuo-tactile stimuli lead to an illusory sense of ownership over the dummy hand. The body-ownership over the dummy body-parts can be extended to the entire body. In Lenggenhager et al. [2007], for instance, the authors showed that a person observing the virtual body being touched while simultaneously the person is also touched caused an out-of-body experience. Interestingly, they found that the stimulation of specific brain regions, precisely the temporo-parietal junction area, could evoke a similar experience [Blanke and Arzy, 2005]. This region is known to work in consonance with multiple body sensations [Blanke et al., 2004]. Despite numerous studies conducted in this regard, a comprehensive understanding of the sense of ownership is still lacking.

Recent phenomenological interpretations provide a concise scheme for understanding the origin of body-ownership[Gallagher and Zahavi, 2007, De Vignemont, 2011, David and Ataria, 2021]. Ataria [2018] summarised the type of feeling–judgement conflict to clarify the various states of body-ownership. Phenomenolrogically, feelings involve closed sensor–motor loops; by contrast, judgements are more abstract reasoning processes. In summary, feelings give a first-person perspective, i.e. a non-conceptual, pre-reflective, and bottom-up process. By contrast, judgement gives a third-person perspective, i.e. observational, reflective, and top-down process. According to his argument, during an RHI, subjects feel the dummy hand as their hand but never admit (or judge) it as their hand. This conflict between feeling (i.e. this is my hand) and judgement (i.e. this is not my hand) causes ambiguous body ownership (ABO) rather than a specific sense of body-ownership [De Vignemont, 2011]. The fact that the results of the two processes are not necessarily consistent motivated us to consider that malleability of body image originates during the mismatched feeling–judgement process.

The free-energy principle (FEP) (or Bayesian inference) is a promising theory for understanding the flexibility of our perceptual system [Friston et al., 2014]. According to the FEP, we can change our beliefs whenever confronted with unexpected situations (i.e. high saliency). The perceptual system updates prior hypotheses with the more preferable posterior hypotheses. The FEP suggests that appropriately adjusting the hypotheses in response to the environment corresponds to the malleability of body image. The RHI is, therefore, related to the corruption of sensorimotor predictive cycles. Phenomenologically, this sensor-motor prediction error indicates a corruption of the feeling system. Most studies do not proceed further than this interpretation of corruption at the feeling level; however, as discussed earlier, the corruption of the feeling system does not provide an adequate explanation for ABO because we must also consider the top-down judgement level.

Tsakiris et al. [2011] investigated the interaction between bottom-up and top-down processes through interoceptive and exteroceptive perceptions [Apps and Tsakiris, 2014, de Vignemont and Alsmith, 2017]. They showed that the intensity of RHI is due to the mismatch between interoceptive and exteroceptive predictions. In their experiments, subjects who predicted their heartbeats more precisely experienced less illusion of ownership during the RHI compared with people who predicted their heartbeats less precisely [Tsakiris et al., 2011]. Based on their results, they speculated the existence of an interdependent relationship between interoceptive and exteroceptive predictions, i.e. high interoceptive saliency suppresses exteroceptive saliency, resulting in ABO (see also Tsakiris [2018]).

A remarkable relationship between RHI subjectivity and FEP was proposed by Tsakiris [2018]; however, FEP per se is fraught with an issue that is open to the following questions. (I) In FEP, judgement is a process for selecting the highest saliency among possible saliencies. As discussed in Ransom et al. [2020], these saliency selections cannot explain human affective behaviour for low saliency. This limitation implies that the saliency-driven explanation is insufficient for human experience. If we accept (I), then we agree that there is no conflict between feeling and judgement because the judgement system mechanically picks up the highest saliency; in this case, the intensity of RHI is only ascribed to the feeling system. If we do not accept (I), then we agree that the sensor–motor cycle holds both feeling and judgement; in this case, we neglect the essential difference between feeling and judgement owing to the compression of two different systems into one sensor–motor system.

Integrated information theory (IIT) is expected to explain human consciousness [Balduzzi and Tononi, 2009, Tononi et al., 2016, Oizumi et al., 2016a,b]. The basic concept of IIT is summarized in the following propositions: the degree of consciousness can be measured as the difference between the entire system and the sum of its parts; that is, the irreducibility of the entire system to its parts is the key to understanding the experience of consciousness [Balduzzi and Tononi, 2009, Tononi et al., 2016]. Notably, IIT also includes feeling and judgement in the perceptual process. Feeling corresponds to the local process and represents the information process of the subsystem (i.e. the subsystems of the system); judgement, in contrast, corresponds to the global process and represents the information process of the entire system. Importantly, IIT rejects any external evaluations as an explanation for consciousness; therefore, there is no superiority between feeling and judgement. In this sense, judgement maintains a parallel relationship with feeling in IIT in contrast to FEP. The difference between feeling and judgement is revealed only by subjective experience; thus, IIT can mathematically evaluate body malleability induced by the conflict between feeling and judgement.

In this study, we examined the relationship between subjective experience and objective bio-signals during an RHI using IIT. Although IIT is usually applied to the complex neural network in the brain, in this study, we picked up seven physiological data from the body (respiration, heartbeat, and skin conductance from body processes) and brain (EEG with electrodes on midline frontal (Fz), vertex (Cz), midline parietal (Pz), and midline occipital (Oz) regions for the brain process). We focused on these data for the following reasons. As highlighted by Tsakiris [2018], the interaction between interoceptive (body) and exteroceptive processes (brain) might explain the degree of ABO during RHI. It is not always necessary to examine the entire brain to estimate a subjective experience. Their result suggests that the degree of conflict between body and brain, rather than that of the entire brain process, is enough to estimate the subject’s experiences during a stimulus.

## 2 The basic concept of IIT

Let us first explain IIT concepts for the sake of readers unfamiliar with IIT. Although many concepts exist in IIT, we only focus on three: minimum information partition (MIP in short), main complexes, and integrated information Φ.

IIT deals with the intrinsic, rather than extrinsic, information; it only depends on inner variables. Typical information theories used in biology focus on the relation between external inputs and their results. In this setting, the system itself becomes a black box. The aim of IIT is to consider this black box in terms of “what the system is” rather than “what the system does” [Albantakis and Tononi, 2015]. The basic concept of IIT expresses mutual information between the past and current system’s states; mathematically, this is expressed as *I*(*X*(*t* − *τ*); *X*(*t*)).

However, this simple mutual information contains redundant information on the system’s integrity, which should be measured as the system’s irreducibility to its parts (subsystems, in IIT). The information obtained from the subsystems must be subtracted from the entire information *I*(*X*(*t* −*τ*); *X*(*t*)). The rest of the information represents the system’s integrity; therefore, the problem for IIT lies in the determination of the system partitions.

This study applies IIT as the mismatch decoding proposed by Oizumi et al. [2016a] because Φ* satisfies the following properties: (i) Non-negativity: the lower bound of Φ* is zero; (ii) sensitivity to noise correlation: Φ* deals with external correlated noise [Mediano et al., 2019]; and (iii) applicability to continuous variables. Sometimes, the IIT as the mismatch decoding is classified as “IIT 2.0” to distinguish with IIT 3.0 [Oizumi et al., 2014, Tononi and Koch, 2015, Albantakis et al., 2019].

### 2.1 Integrated information (Φ*) and minimum information partition (MIP)

As mentioned in the previous section, the mode of partitioning is essential to determining the system’s integrity. The MIP is the system partition where the system’s integrity is the minimum. The system’s integrity in Φ* is given by

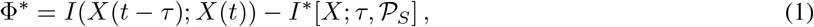

where *I*(*X*(*t* −*τ*); *X*(*t*)) is the mutual information between the current *X*(*t*) and past states *X*(*t* −*τ*); *S* is the set of all nodes of a given system; 𝒫_*S*_ is the set of all bi-partitions (total 2^|*S*|^ −1 partitions); *I*\*(*X*; *τ,* 𝒫_*S*_) is a “hypothetical” mutual information, indicating the mismatched decoding in the partitioned probability distribution. Precisely, *I*\*(*X*; *τ,* 𝒫_*S*_) is given as the partition max_*β*_ *Ĩ*(*β*; *X, τ*, 𝒫_*S*_) that minimizes Φ* (see [Oizumi et al., 2016a, Mediano et al., 2019] for further details about this expression).

Because Φ* depends on the partition *π*(∈𝒫_*S*_), the MIP is the partition that minimizes the integrated information.

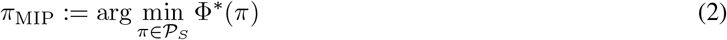

The integrated information of *π*_MIP_ is expressed as Φ*(*π*_MIP_). We simply denote Φ*(*π*_MIP_) as Φ_MIP_. Notably, if Φ_MIP_ equals zero, the parts of the system are mutually independent; that is, there is no interaction between the parts. In this sense, Φ_MIP_ characterises the system’s irreducibility to its parts.

### 2.2 Main complexes

Generally, we can compute Φ_MIP_ for any subsystem in the system, not only for the set *S*. We denote each Φ_MIP_ for a subsystem *T* as 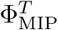, where *T* ⊂ *S*. A ‘complex’ is a subsystem *C*(⊂ *S*), where 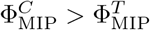 for all supersets of *S*. From this definition, we can define the main complexes as those with the local maximum 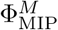.

#### Definition (Main complex)

A main complex is a complex *M* satisfying 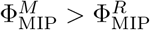 for any subsystems *R* ⊂ *M*. The definition says that if two main complexes exist (say, *A* and *B*), then they are exclusive. IIT researchers consider these main complexes to be the system’s information core that might be related to our conscious experience. The validity of this assumption remains to be demonstrated, although there is no doubt that such information cores play a vital role in living systems [Marshall et al., 2017, Sheneman et al., 2019, Niizato et al., 2020].

## 3 Application to RHI

### 3.1 Participants

Twenty-two healthy male adults (mean age, 21.1 years; SD, 0.5) were recruited among the students at Nagaoka University of Technology. All participants gave written consent to participate after receiving an explanation of the procedures involved. The study was conducted according to the principles of the Declaration of Helsinki and was approved by the Ethics Committee of Nagaoka University of Technology.

### 3.2 Procedure

Participants sat on a chair with their left arm on a table and their palms facing down. The experimenter stood opposite the participant across the table. A realistic-looking left rubber hand was placed to the right of the participants’ real left hand. The distance between the index finger of the rubber hand and the index finger of the real hand was 15 cm. Two grey towels covered the space from the participants’ shoulder to the participants’ wrist and the proximal end of the artificial hand. A 45 cm × 60 cm standing screen between the rubber and real hand prevented participants from viewing their own left hand (S1 Fig). Two experimental conditions were performed in our study: synchronous and asynchronous stroking condition. The experiments comprised two blocks corresponding to these conditions. The sequence of the blocks was randomized and counterbalanced across participants. Each block lasted for 15 min with a few minutes of inter-block intervals: 5-min rest periods (Rest Phase: Pre and Post) before and after a 5-min stroking period (Stimulus Phase). The rubber hand and unseen hand were stroked by two brushes either synchronously or asynchronously during the stroking period. The trained experimenter indicated the predetermined pattern and frequency (40 bpm) of stimulation using a metronome through an earphone. In the asynchronous condition, the dummy hand was touched immediately after touching the hidden real hand by the metronome; that is, every visual stimulus followed a tactile stimulus in the asynchronous condition. Therefore, the tactile stimulation was identical for both conditions. After the second rest period, the participants answered a questionnaire about their subjective experiences during the stroking period.

### 3.3 Measurement

#### Physiological Measurement

Throughout each experimental block, electroencephalogram (EEG), electrocardiogram (ECG), respiration air flow (RESP), and electrodermal activity (EDA) were recorded using a bio-amplifier (Polymate Pro MP 6000, Miyuki Giken Corp., Tokyo, Japan) at a sampling rate of 250 Hz and with a 24-bit resolution, using a 50-Hz notch filter. Especially, for the EEG, four signals (Fz, Cz, Pz, Oz) were measured at the locations of the four electrodes on the scalp according to the international 10-20 systems for measuring the four EEG signals (Fz, Cz, Pz, Oz); here, we used a high-frequency filter of 50 Hz and a low-frequency filter of 1 Hz.

#### Questionnaire

The questionnaire included nine statements from the original RHI study [Botvinick and Cohen, 1998] presented on a computer screen in a random order S1 Table. Participants indicated their response by clicking on a visual-analogue scale ranging from strong agreement (100) to strong disagreement (0) including neither agreement nor disagreement (50). All the questions and instructions were presented in Japanese. Questionnaire data from one participant were excluded from the analysis because they were lost owing to mechanical issues.

### 3.4 Data setting for IIT 2.0 application

Kanai and Oizumi et al. proposed approximation methods for the computational problem of IIT [Oizumi et al., 2016b,a, Kitazono et al., 2018, Hidaka and Oizumi, 2018]. This study applied their “Practical Φ Toolbox for MATLAB” to the physiological data; we considered only seven physiological data for the 22 subjects. The exhaustive method (computing all possible MIPs) can be applied in realistic computation time for such a small system.

Another parameter is time delay *τ* (IIT has only two constraints: partitions *π* and time delay *τ*). To choose the suitable parameter, we computed Φ_MIP_ for *τ* from 0 to 250 frames. The time delay *τ* is the point where the mean is maximum Φ_MIP_ for all subjects. We found that this *τ* is approximately 50 frames (i.e. 0.2 s). This value seems to be suitable because the value is the same as the human response time (S2 Fig).

The time series of each raw data was 225000 frames (250 frames per second). Because we applied a 50 frame-delay (0.2 s) for the computation of IIT 2.0, sufficient data series for IIT 2.0 are required. We attempted 500 frames (2 s), 750 frames (3 s) and 1250 frames (5 s) for IIT 2.0 computations — the time window shifts 250 frames each. The first blank data for some frames (500, 750, 1250 frames) were set at zero (only the first five steps at most for 1250 frames), which sum up to 900 data sets for each subject. The results for all frame samples were almost the same (S2 Table).

We extracted all measures related to integrated information for 900 steps (1 s for one step). The first third (from 0 − 300 steps) and the last third (from 600 − 900 steps) correspond to the pre- and post-stimulus resting states. The middle section (from 300 − 600 steps) is where the synchronous or asynchronous stimulus was applied. For all statistical tests, we detrended the obtained Φ time series because our focus was on the variation due to the stimulus for our rubber hand experiments. A negative value of Φ_MIP_ is due to this treatment.

## 4 Results

### 4.1 MIP (minimum information partition)

We examined the average tendency of MIP during the experiments. As earlier discussed, the MIP gives two measures for estimating the system’s state: (i) MIP cut, which divides the two subsystems by the weakest information flow; (ii) Φ_MIP_, which is the degree of information loss by the MIP cut. Φ_MIP_ is positive by definition, otherwise (i.e., if it were equal to 0), the two subsystems would be completely independent systems.

This study used seven data sets (i.e. respiration (Res), heartbeat (ECG), and skin conductance (EDA) for BODY; midline frontal (Fz), vertex (Cz), midline parietal (Pz), and midline occipital (Oz) region for BRAIN as EEG) for IIT analysis. We considered that these body–brain signals work as a system (*S* ={Res, ECG, EDA, Fz, Cz, Pz, Oz}). If the rubber hand stimulus causes a difference in the system, the relation among these seven data sets will change. This difference also reflects the MIP information.

Figure 1A shows the time series of the mean Φ_MIP_ for all 22 subjects. The Φ_MIP_ drops at the stimulus phase for both cases (Figure 1B). Decreasing Φ_MIP_ values suggests that connections between subsystems become weaker than pre- and post-stimulus. This result is interesting because similar changes of Φ_MIP_ occur in both of SYNC and ASYNC conditions.

**Figure 1:**
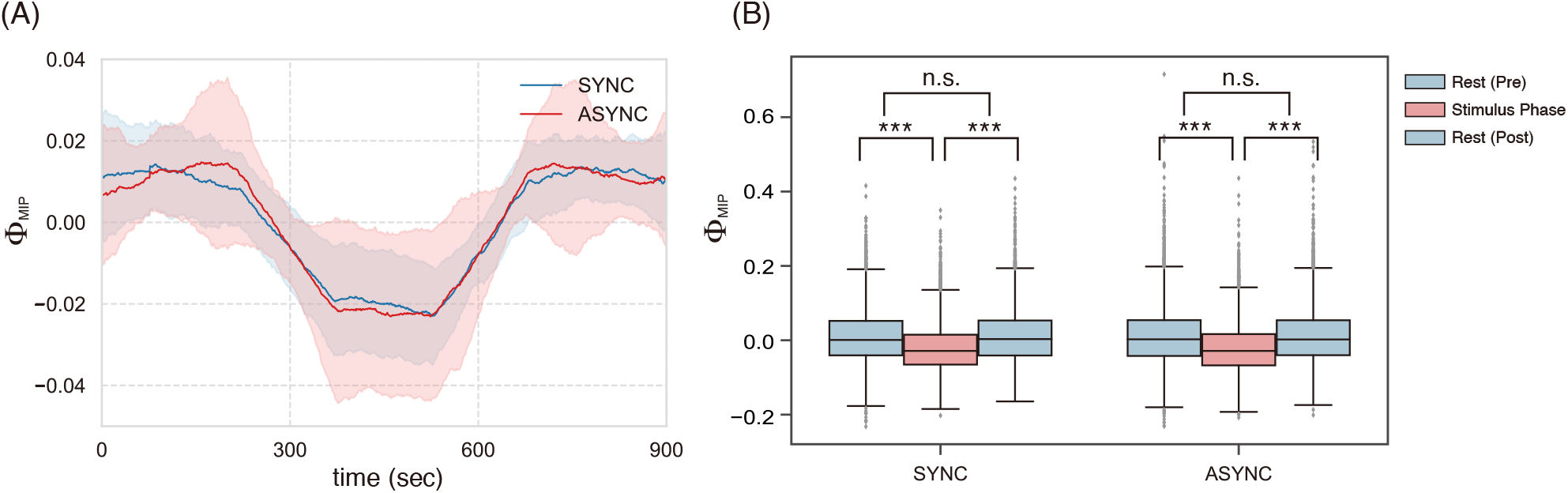
(A) Time series of mean Φ_MIP_ for both conditions (blue: SYNC, red: ASYNC). The stimulus phase is during 300^600 s. The other regions indicate resting states (no stimulus). The computational time frame for IIT is 500 (2 s). Moving-average (sample length 150 steps) was performed on the time-series for attaining easy to see temporal changes of values. (B) Φ_MIP_ of SYNC and ASYNC values drop significantly compared with each rest states (\**p* < 0.05,* * *p* < 0.01, * * \**p* < 0.005).

Next, we examine the details of the MIP cut. The most remarkable tendency is the cut between {Oz} and {Res, ECG, EDA, Fz, Cz, Pz}. More than 50% of MIP cuts were of this type (Figure 2A, B); furthermore, during the stimulus phase, the frequency of this cut increased statistically. Although their frequency increased, the Φ_MIP_ of {Oz}-cut decreased. The implication of this tendency is suggestive. It is a well-known fact that the occipital region (Oz) is involved in the visual information process. The high frequency of {Oz}-cut reflects the discrepancy between visual information and tactile information. The low Φ_MIP_ during the stimulus means the visual process in Oz separates from other information processes (Figure 2C).

**Figure 2:**
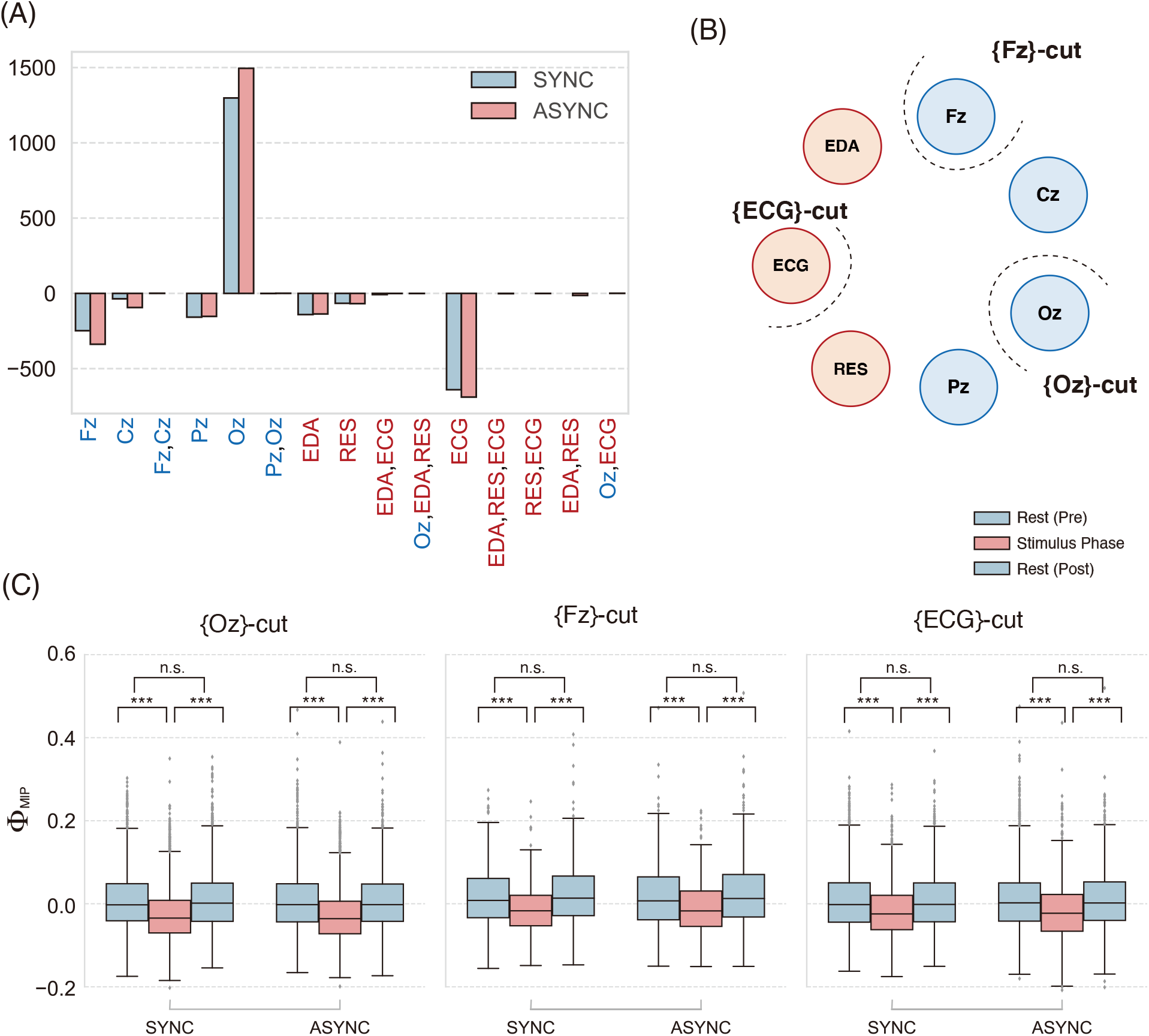
(A) Difference in the *π*_MIP_ frequency for pre- and intra -stimulus: Fr(*π*_MIP_ |SP) −Fr(*π*_MIP_ |PRE). The computational time frame for IIT is 500 (2 s). The positive values indicate that a certain type of *π*_MIP_ increases during the stimulus. Blue (red) bars show the frequency differences in the synchronous (asynchronous) condition. Blue (red) letters show data from the brain (body) system. (B) Image of MIP-cut location for the system *S*. {Oz}-, {Fz}-, {ECG}-cut are subsets with three largest differences shown in (A). (C) Φ_MIP_ for each MIP-cut location. For all cases, the Φ_MIP_ drops during the stimulus (\**p* < 0.05, * * *p* < 0.01, * * \**p* < 0.005). No after-effects were observed.

For the remaining patterns, the MIP cut frequencies decreased after the stimulus was applied (Figure 2A, B). The decreased frequency of the rest compensates for the increased frequency of {Oz}-cut. Interestingly, in the first case we observed, the cut was rarely located between the body and brain systems. For almost all cases, the cut located the brain system. This fact suggests that the body and the brain system is not the weakest link. The second most frequent Φ_MIP_ cut locates between {ECG} and {Res, EDA, Fz, Cz, Pz, Oz}. Taking account of this cut during stimulus, the heartbeat (ECG) merges into the cycle of the brain–body system during the stimulus. Even if {ECG} was selected as the cut, its integrity decreased during the stimulus. This aspect resonates with Tsakiri’s result [Tsakiris et al., 2011].

To end this subsection, we highlight the following fact: Φ_MIP_ for SYNC and ANSYC condition during RHI shows the statistical difference between the two conditions observed (Welch’s t test: *df* =13159, t-stat= 2.2561, *p* < 0.05). This result seems interesting because half of the subjects in our study reported a change of body ownership during both conditions. Many of the individual differences were not in nature but in degree. This logic also applies to the next subsection in **Main complex analysis**.

### 4.2 Main complexes analysis

We examine the average tendency of the main complexes for SYNC and ASYNC condition. The concept of the main complex accompanies some other concepts; namely, (i) 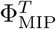, which is the degree of information loss by partitioning the system into independent subsystems. The manner of partitioning system *T* is determined by MIP: 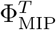; (ii) Complex, where all subsystems satisfy 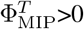 and 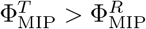for any superset *R* of *T* ; and (iii) Main complex, where the complex *M* satisfies 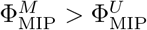 for any subset *U* of *M*.

Because the Φ_MIP_ represents the irreducibility to the system’s parts, the high Φ_MIP_ values measure the system’s integrity. Notably, the system may include several main complexes. For a subsystem to be the main complex, it is sufficient that it does not contain other complexes (Lemma 4 in Kitazono et al. [2020]). In our analysis, the body–brain system has up to ten or more main complexes. The integrity of the system can be estimated as the sum of 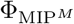 of all main complexes *M* in *S* (denote 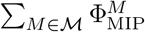 where ℳ is a set of main complexes of system *S* ={Res, ECG, EDA, Fz, Cz, Pz, Oz}).

Figure 3A shows that 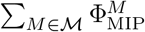 increases only in the stimulus phase. In contrast to the decreasing 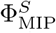 for the entire system *S*, the integrities of the particular subsystems increased (Figure 3B). While the entire system splits up into relative independent parts, specific subsystems compensate by their integrity. This tendency can also be observed in the number of complexes. The number of irreducible subsystems (parts) increased during the stimulus phase (S4 Fig). Furthermore, in contrast to the 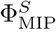, an after-effect was observed in the main complex. This means that fluctuations in the system integrities remained even after the stimulus had ended.

**Figure 3:**
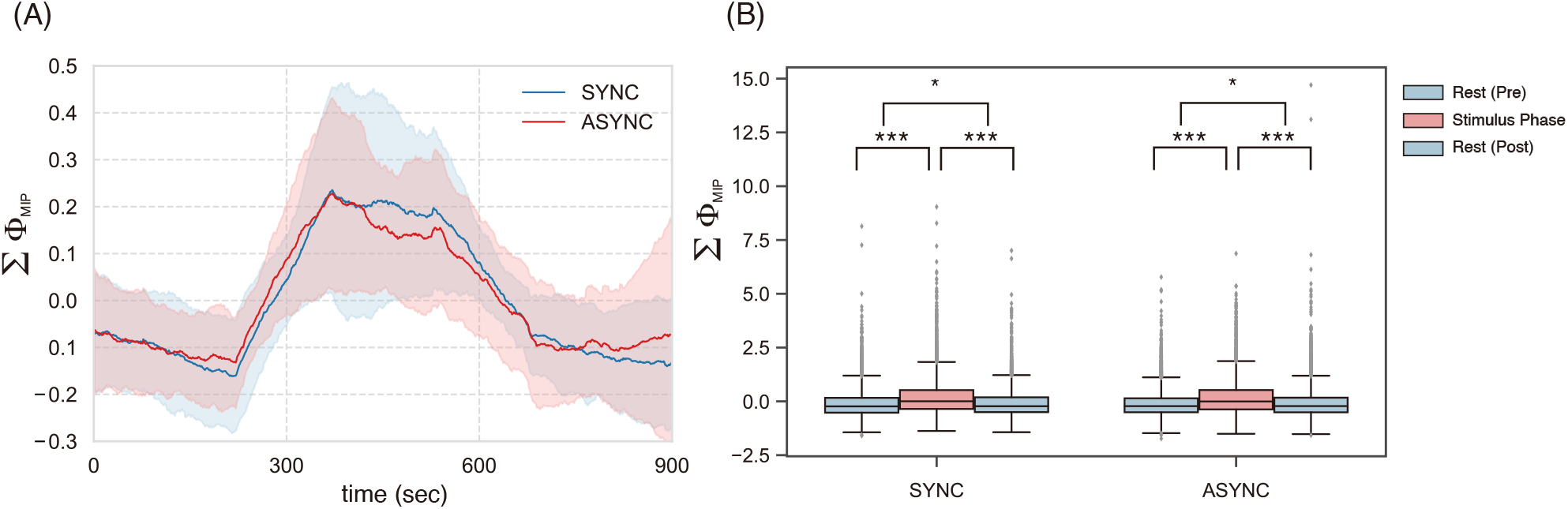
(A) Time series of mean is 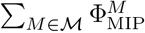 for both conditions (blue: SYNC, red: ASYNC). The stimulus phase is 300^600 s. The other regions are resting phase (no stimulus). The computational time frame for IIT is 500 (2 s). Notably, the moving-average (sample length 150 steps) was performed on the time-series for attaining easy to see temporal changes of values. (B) 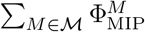 of SYNC and ASYNC values rises significantly compared with each rest phase (\**p* < 0.05, * * *p* < 0.01, * * \**p* < 0.005). In contrast to MIP analysis, an after-effect was observed for both conditions.

The content of the most vital main complex (i.e. max 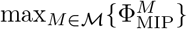: the main complex *M* with the highest 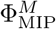) is more suggestive. Figure 4A shows that the frequency of the main complex containing all elements (or 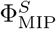) dropped during the stimulus phase. Here, the value of 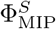 indicates that the body–brain unit itself is irreducible to its parts. Metaphorically, the oneness of the body–brain system was corrupted during the stimulus. After the stimulus phase, the frequency of this main complex recovered.

**Figure 4:**
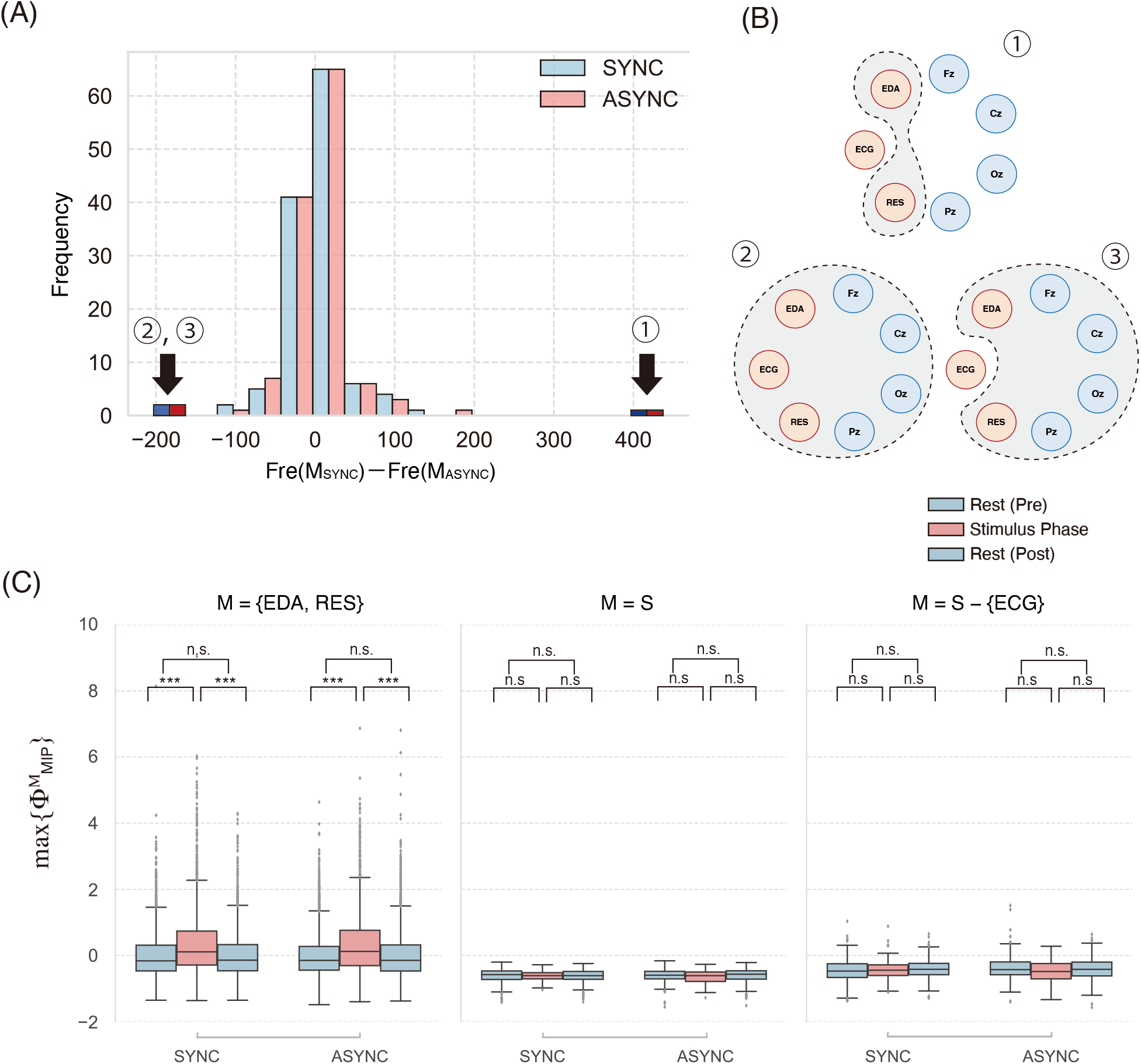
(A) Histogram of the difference of the 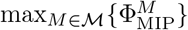 frequency for pre- and intra-stimulus: Fr(*M* ∈ℳ| SP) −Fr(*M* ∈ℳ |PRE). Because many main complexes exist, we only analyse the outliers (indexed 1, 2, 3). Blue (red) bars show the frequency in the synchronous (asynchronous) condition. (B) Image of the main complex for the system *S*. {EDA, RES}, *S, S*-{ECG} are three highest differences. (C) 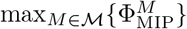 for each main complex. Only *M* ={EDA, RES}, the Φ_MIP_ rises significantly during the stimulus. The other two values are negative throughout three phases. (\**p* < 0.05, * * *p* < 0.01, * * \**p* < 0.005).

The other main complexes compensated for the decreasing frequencies of the complexes *S* (Figure 4B, C). The main complex {EDA, RES} is the most significant: skin conductance and respiration. The 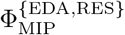 also increased significantly during the stimulus (Figure 4C). The integrity only involves the body parts. Another high frequent main complex is {Fz, Cz, EDA, RES}, containing {Fz, Cz} pairs. The frequency of this complex increased during the stimulus (the 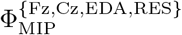 also increased: Welch’s t test: *df* =553, t-stat=-4.0359, *p* < 10^−6^). This might relate to how the subject attempts to match past information and current information.

As was observed in MIP analysis, 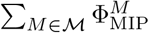 for the SYNC condition during the stimulus is statistically more significant than that of the ASYNC (Welch’s t test: *df* =13177, t-stat=-2.3734, *p* < 0.05). Although both conditions changed the subject’s body ownership, the Φ value reflected the difference between the two subjective reports of the participants.

### 4.3 IIT 2.0 estimates the degree of ownership

In this final subsection, we discuss how IIT 2.0 can estimate subjective reports of body ownership. The original purpose of IIT was to measure the degree of consciousness theoretically. Although the estimation of consciousness itself is still beyond our computational power, this aim could be attained through our small body–brain system.

We have thus far observed the average effects of (a-)synchronous stimulus on each brain–body system; however, as we have seen, there are individual differences in their effects (S3 Fig). While some subjects had a strong ownership illusion for their hand during the stimulus, some subjects did not. Some subjects had an ownership illusion for both conditions, while others only had it for the rubber hand condition. Although, statistically, SYNC subjects felt more intense ownership illusion compared with ASYNC subjects, it is desirable to know why such individual differences emerge.

We averaged the subjective values for Q1–Q3 questions, which indicate ABO for each subject, for both conditions to estimate the subjective reports. This simple computation suggests that the SYNC condition has a strong ownership illusion compared with the ASYNC condition. We use these mean values for each subject for the following analysis.

We begin by answering the following question: can the integrated information and Φ* for each subject account for such subjective individual differences? The answer is negative. For both conditions,∑ Φ_MIP_ (abbr. 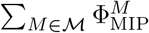) shows low correlations (SYNC: Pearson’s correlation test, *n* = 21, *r* = 0.28, *p* = 0.21. ASYNC: *n* = 21 *r* = 0.20 *p* = 0.38). Interestingly, ∑Φ_MIP_ itself does not reflect the subjective reports. In our system, at least, the ∑Φ_MIP_ is not the proper estimator of subjective reports.

We examine the difference in ownership intensities between the SYNC and the ASYNC condition. After subtracting the baseline (mean pre-stimulus phase for each subject), we take the difference of stimulus phase between the SYNC and the ASYNC. We define the difference in ABO intensities as the sum of this time series. Mathematically, this measure can be expressed for each subject indexed *i*:

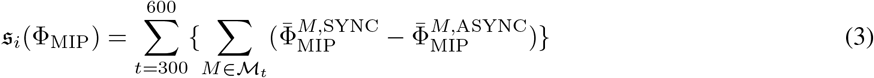

where 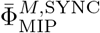 means that the integrity of main complex *M* at time *t* for the SYNC condition subtracted the baseline. Figure 5 shows the correlation between the subjective difference and ownership intensities (Pearson’s correlation test: *n* = 21, *r* = 0.56, *p* < 0.005). The gap between the two experimental settings can comprehend the subjective difference for their ABO. Our result shows that the Φ_MIP_ gaps, rather than of Φ_MIP_ itself, can estimate the subjective individual differences for the different experimental settings.

**Figure 5:**
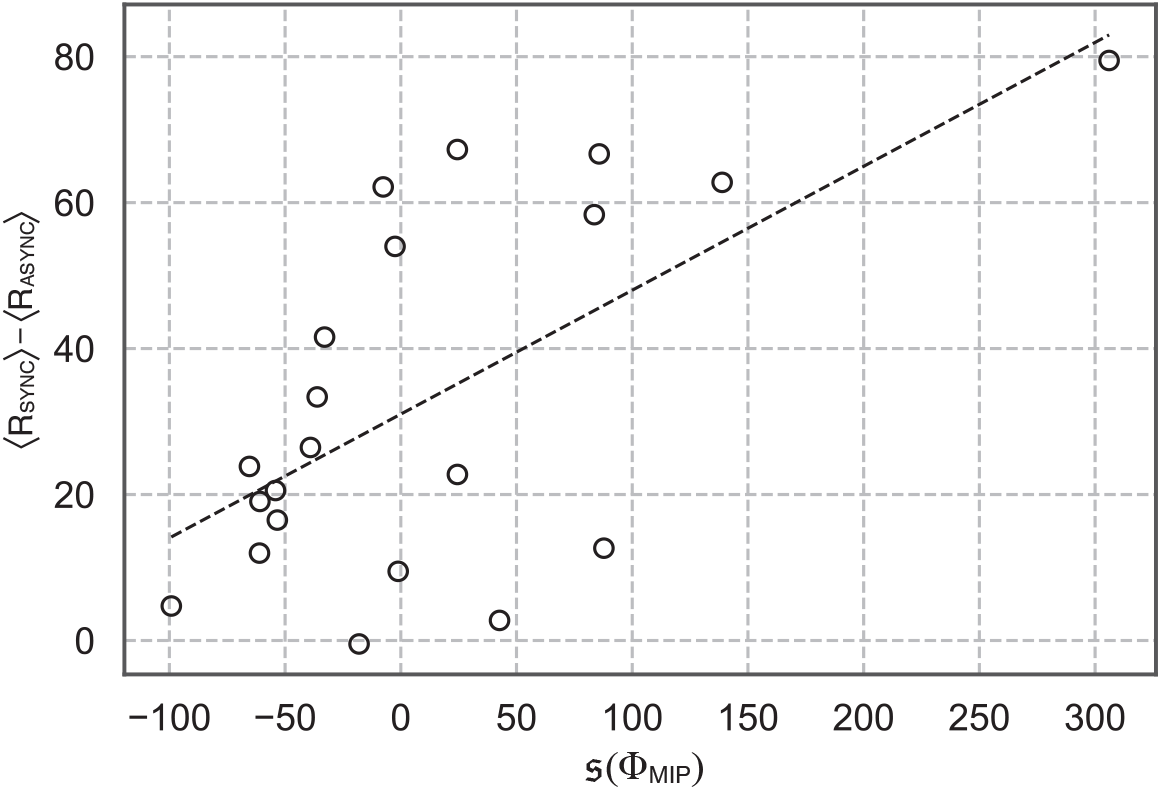
Relationship between the difference of mean subjective report ⟨ *R*_SYNC_⟩ − ⟨*R*_ASYNC_⟩ and 𝔰 (Φ_MIP_) (see Eq. 3) for each subject. The correlation efficient is *r* = 0.56 (Pearson’s correlation test: *n* = 21, *p* < 0.005)

## 5 Discussion

Throughout this study, we have examined two aspects of ABO induced by the rubber hand stimulus: the average tendency and the individual difference.

### 5.1 Average tendencies for (a-)synchronous stimulus of the rubber hand

Applying IIT 2.0 to the small body–brain system, we found some new insights. The minimum information partition (MIP) reveals that the system’s independence emerges during the stimulus for both experimental conditions. The location of the MIP cut indicates that the disjunctive parts are not body–brain pairs but intra-brain or intra-body pairs. The main complex analysis gives this trend a more precise picture – the frequency of all-elements main complex *S* decreases during the stimulus phase. Before the stimulus, the body–brain system works as one irreducible system. Although the main complex of all elements *S* exists even in the stimulus phase, the degree of this irreducibility 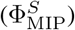. decreases. These results agree with our intuition on these experimental settings.

The main complex shows high integrities (i.e.∑ Φ_MIP_) during the stimulus on average. The disconnected system in terms of MIP compensates for its disconnection by increasing the integrity of its parts. The high irreducible parts, for example, contain Fz, Cz, Res, ECG. Considering that MIP locates at Oz, the complexes, including Fz and Cz, also have a compensated meaning. In the stimulus phase, the brain might raise its integrity for the working memory (Fz) and motor function (Cz) instead of separating the inconsistent visual information (Oz). This relation makes sense if we consider that the motor-perception with the subject’s short term memory (including multisensory integration at the ventral premotor cortex [Ehrsson et al., 2004, Gentile et al., 2015]) attempts to counterbalance the mismatch in the visual system. The body integrity (Res and EDA) is also observed in the stimulus phase. Notably ECG (i.e. heartbeat) is excluded from this body-integrity (the MIP cut locates at {ECG} in Figure 2B). This relative independency of the heartbeat might be related to Tsakiris et al. [2011] results.

Although there were some differences between SYNC and ASYNC conditions, we also observed considerable similarities in Φ_MIP_ variations. This similarity suggests that the same physiological phenomena might occur in both SYNC and ASYNC conditions. Interestingly, Perez-Marcos et al. [2018] pointed out that body image distortion elicits can be elicited by ASYNC condition, but not SYNC condition. Their results imply that the ASYNC condition also changes latent body cognition, and our analysis can associate with their results.

### 5.2 Individual differences of ABO with IIT 2.0

Although our average IIT analysis succeeded in explaining the general tendencies, individual differences still exist. Some subjects experience more ABO than ASYNC condition; others experience ABO as the same as the same condition. Previous studies try to explain these various individual differences from the physiological perspective [Armel and Ramachandran, 2003]; however, there are no consistent conclusions.

We found that the ∑Φ_MIP_ value itself does not reflect subjective reports. Instead, the difference between them correlates the subjective reports. This result suggests that subjective evaluation is not determined by a single experience but rather by comparing different experiences. We also attempted the same analysis applying the peak–end rule, which states that how they felt at its peak and at its end determines judgement on a given experience [Kahneman et al., 1993]. The result was the same.

Why does the difference in Φ_MIP_ correlate with individual subjective report? As discussed in our result, this might be due to the manner of reporting. The subject was required to answer their feelings of ABO from 0 −100^°^. Almost all subjects would have been asked this question for the first time. This setting means that the subject cannot access his or her past ownership experience. The meaning of ownership attains its definitive status after comparing the different types of ownership. Without such different experiences, the subjective report has no reference point. Therefore, the anchoring effect of the subjective report might be cancelled out by considering differences between two different ownership experience.

### 5.3 Evaluate subjective experience: free energy and integrated information

Sometimes, the rubber hand illusion has been interpreted in the context of Bayesian inference [Samad et al., 2015, Rood et al., 2020]. Bayesian inference says that the subject attempts to eliminate perceptual inconsistency by replacing it with a new hypothesis. This assumption agrees with the free energy principle (FEP) proposed by Friston. Because perceptual mismatch increases the system’s free energy (or surprise), it attempts to decrease it with an updated hypothesis.

How does this FEP relate to IIT? A possible answer is that the relation between FEP and IIT is that they are ‘two sides of the same coin’. Some simulation studies help us with this understanding. Although there are many simulation studies on IIT, almost all of them make the same observation; that is, the integrated information mainly increases in the learning phase [Albantakis et al., 2014, Abrego and Zaikin, 2019, Sheneman et al., 2019]. Once the order of the system has been established, the Φ values decrease [Sheneman et al., 2019].

If we observe that the Φ originally refers to consciousness, these tendencies suggest much awareness needed during the learning phase. However, Solms and Friston [2018] have indicated that reducing “surprise” means losing the conscious process (establishing the automatic response to the external environment [Solms, 2019]). In this sense, we might say that the IIT and FEP observe the same object from different perspectives.

In this context, the RH stimulus might trigger a learning phase from the IIT perspective. From the point of view of the FEP, this also refers to the generation of a system’s “surprise”; however, the process is not straightforward. As earlier noted, FEP hardly explains the conflict between feeling and judgement on sense of ownership by itself; IIT can access this conflict. Furthermore, our IIT analysis shows that the Φ value in IIT also fails to capture subjective experiences.

How can we relate our results to the subjective reports? Our small system cannot explain that consciousness itself might be one of the possible interpretations. Nevertheless, the most plausible interpretation lies in the process of subjective feeling estimation—, that is, the subject is required to report their feelings after each experiment. In this setting, the subject reports their judgement of experience rather than their ownership experience themselves. Judgement is not the direct experience, but an indirect experience mediated by language. The subjective report does not reflect subjective experiences directly (note that this judgement differs from the one we used in the feeling-judgement conflict; rather, it is the second-order judgement for this conflict).

There are many criticisms of IIT. One of the main criticisms stems from the statement that Φ can measure raw consciousness [Cerullo, 2015, Lombardi and López, 2018]. Indeed, this type of statement contains complex philosophical issues, which are not easy to solve. However, if we consider IIT as a measure of subjective experience on the second-order judgement level, then our study suggests that IIT may provide the mathematical framework for phenomenological descriptions of subjective experience.

## Supporting information

**S1 Fig. Experimental set-up in the stimulus phase. A dummy left hand was placed in front of the participants. The participants’ left hand was hidden out of view. The right hand with a probe to measure EDA rested on a table. EEG electrodes were installed on participants’ scalp. In addition, participants were equipped with ECG electrodes and nostril cannula to measure their respiration flow**.

**S2 Fig**. Φ_MIP_ **values along time delay** *τ* **s for the entire system** *S*. **The frame size is 900 s**. Φ_MIP_ **peak locates at 0.2 s**.

**S3 Fig. Boxplot of rating scores for each questionnaire item. The data is available on S1 Table**.

**S4 Fig. Boxplot of the number of complexes for each main complex. The tendency is the same as the** 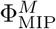(*M* ∈ ℳ)**distribution in Figure 4**.

**S1 Table. Questionnaire items [Botvinick and Cohen**, **1998]**.

**S2 Table. All statistical tests for each condition**.

**Table 1:**
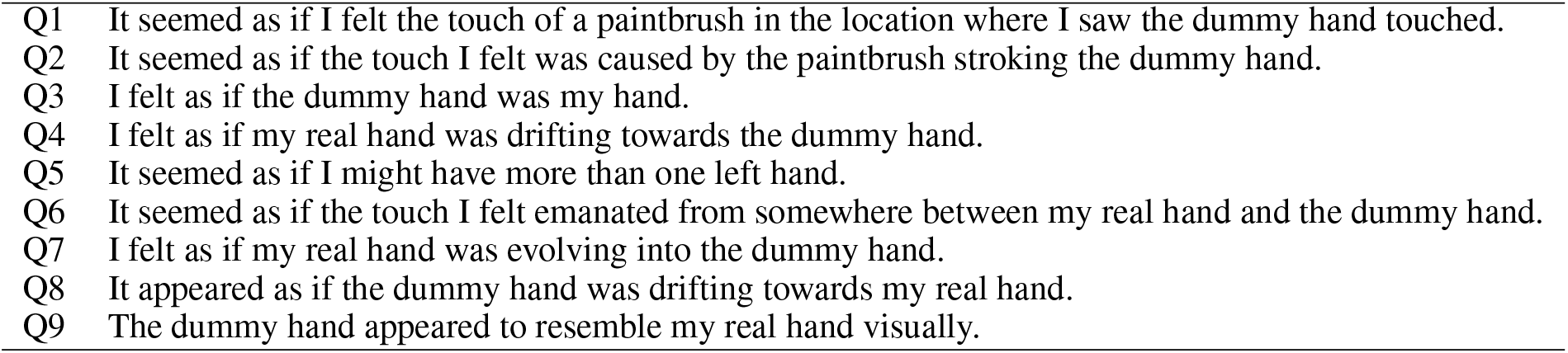
Questionnaire items [Botvinick and Cohen, 1998].

